# Changes in physicochemical properties and microbial community succession during leaf stacking fermentation

**DOI:** 10.1101/2023.02.06.527411

**Authors:** Guanghai Zhang, Lu Zhao, Wei Li, Heng Yao, Canhua Lu, Gaokun Zhao, Ziyi Liu, Yuping Wu, Wanlong Yang, Yongping Li, Guanghui Kong

## Abstract

Leaf stacking fermentation involves enzymatic actions of many microorganisms and is an efficient and environmentally benign process for degrading macromolecular organic compounds. We investigated the dynamics of metabolite profiles, bacterial and fungal communities and their interactions during fermentation using cigar leaves from three geographic regions. The results showed that the contents of total sugar, reducing sugar, starch, cellulose, lignin, pectin, polyphenol and protein in cigar tobacco leaves was significantly decreased during fermentation. Notably, the furfural, neophytadiene, pyridine, benzyl alcohol, geranyl acetone, 2-butanone, 3-hydroxy-, N-hexanal, isoamylalcohol and 2,3-pentanedione were important features volatile aroma compounds during fermentation. The α-diversity of fungi and bacteria initially increased and then decreased during fermentation. The microbial diversity was influenced by fermentation stages and growing locations, in which the fermentation stages had greater impacts on the microbial diversity than locations. Microbiome profiling had identified several core bacteria including *Sphingomonas, Bacillus, Staphylococcus, Pseudomonas, Ralstonia, Massilia* and *Fibrobacter*. Fungal biomarkers included *Aspergillus, Penicillium, Fusarium, Cladosporium* and *Trichomonascus*. Interestingly, the molecular ecological networks showed that the core taxa had significant correlations with metabolic enzymes and physicochemical properties; bacteria and fungi jointly participated in the carbohydrate and nitrogen compound degrading and volatile aroma compound chemosynthesis processes during fermentation. These studies provide insights into the coupling of material conversion and microbial community succession during leaf fermentation.

**IMPORTANCE:** Cigar tobacco leaves are still crude after air curing. Fermentation can eliminate the pungent and bitter taste of the leaves and promote the accumulations of aromatic compounds. Hitherto, there has been no systematic study on the material conversion, metabolic enzymes and microbial community changes and their interactions in fermented leaves. The significance of our research is in identifying the potential metabolic cooperation related to microbial community, which will greatly enhance our understanding of microorganism to convert macromolecular organic to small molecules during the leaf stacking fermentation process.

---

Cigar tobacco leaves are still crude after air curing. Changes in chemical composition that occur during air-curing constitute preparatory rather than the decisive steps on the way from the green tobacco leaves to the satisfactory and industrially acceptable smoking quality (1). It is necessary to carry out sweating or aging for a certain period of time to eliminate the pungent and bitter taste of the leaves and to boost accumulations of aromatic compounds that may originate from the products burned and distilled during smoking (2). Different complex chemical reactions are involved in tobacco aroma formation, including degradation of carbohydrates, degradation of chlorogenic acid, degradation of proteins, Maillard reaction, Strecker degradation and caramelization reactions (3). Tobacco fermentation is a process of biochemical transformation of various organic compounds in tobacco leaves under the synergistic action of inorganic elements, enzymes and microorganisms (4). Chemical reaction effects mainly include redox reactions and Maillard reactions. Organic matter is oxidized by oxygen in the air during the catalysis of inorganic elements (such as iron, and magnesium) (5). There are many enzymes in tobacco leaf cells, which are the main catalytic factors involved in many chemical transformation pathways during tobacco leaf fermentation (6, 7). Frankenburg proposed that tobacco leaf fermentation was a process of substance transformation catalyzed by enzymes (1, 2). The enzyme activity in tobacco leaf cells was higher under high temperature and high humidity conditions. A series of studies has focused on the structure, function and quality improvement of the microbial community before and after cigar tobacco fermentation. There are different kinds of microorganisms on the surface of cigar tobacco leaves from different places. The main microbial species include fungi and bacteria of *Bacillus, Staphylococcus, Corynebacterium, Lactobacillus, Pseudomonas, Penicillium, Aspergillus, Rhizopus* and *Mucormycosis* (8-11). During fermentation, a variety of microorganisms on the surface of tobacco leaves, and cigar tobacco leaves contain organic matter such as sugar, protein and fat needed for microbial propagation and growth. Under suitable environmental conditions, microorganisms multiply and grow and secrete metabolically related enzymes to breakdown and utilize the carbohydrates and nitrogen compounds in tobacco leaves and further degrade them into small molecule metabolites, key to achieving a better and pleasant taste during smoking(4, 12). Microbial communities showed significant correlations with protein, lignin, and cellulose(13). However, the diversity and dynamics of the microbial community and physicochemical properties at different fermentation stages remain unclear. Microorganisms of cigar tobacco leaf change dynamically with space and time. However, previous studies have focused only on the postfermentation stages, and little is known about the diversity and dynamics at different fermentation stages. The current study captured temporal changes in microorganisms at different fermentation stages, thereby assisting in the understanding of the microbial community associated with cigar tobacco leaf fermentation.

After proper fermentation, the smoking quality, leaf appearance and physical properties of cigar tobacco leaves were obviously improved, which was closely related to the chemical changes in tobacco tissues during fermentation. The odor, color, texture, water retention and pH change in a characteristic way. Following air curing, cigar tobacco leaves undergo a fermentation process consisting of material transformation, degradation, acidification, volatilization, etc (14). Hitherto, there has been no systematic study on the quality, enzyme and microbial changes of cigar tobacco leaves during fermentation. This study systematically analyzed the chemical composition, physical properties, metabolic enzymes and microbial community changes and their interactions during the fermentation of cigar tobacco leaves. The fundamental purpose of this study is to reveal the dynamics of material conversion, the composition in bacterial and fungal communities, and the interactions between chemicals and microorganisms.

## RESULTS

### Chemical composition changes during fermentation

The conventional chemical components of cigar tobacco leaf from three locations were analyzed before fermentation(F0) and at different fermentation stages (F1, F2, F3, F4, F5). The results showed that the total sugar (TS), reducing sugar (RS), starch (SH), acid cellulose (AC), acid lignin (AL), pectin (PT), total polyphenols (TP) and protein (PN) in cigar tobacco leaves were significantly decreased during fermentation (Fig. 2A, Fig.S1). There were no significant changes in the contents of total nitrogen (TN), nicotine (NT), potassium oxide (K), chloride ion (Cl), magnesium (Mg), petroleum ether extract (PEE), total amino acids (TAA) and the pH value (Fig. S1).

**FIG 1.**
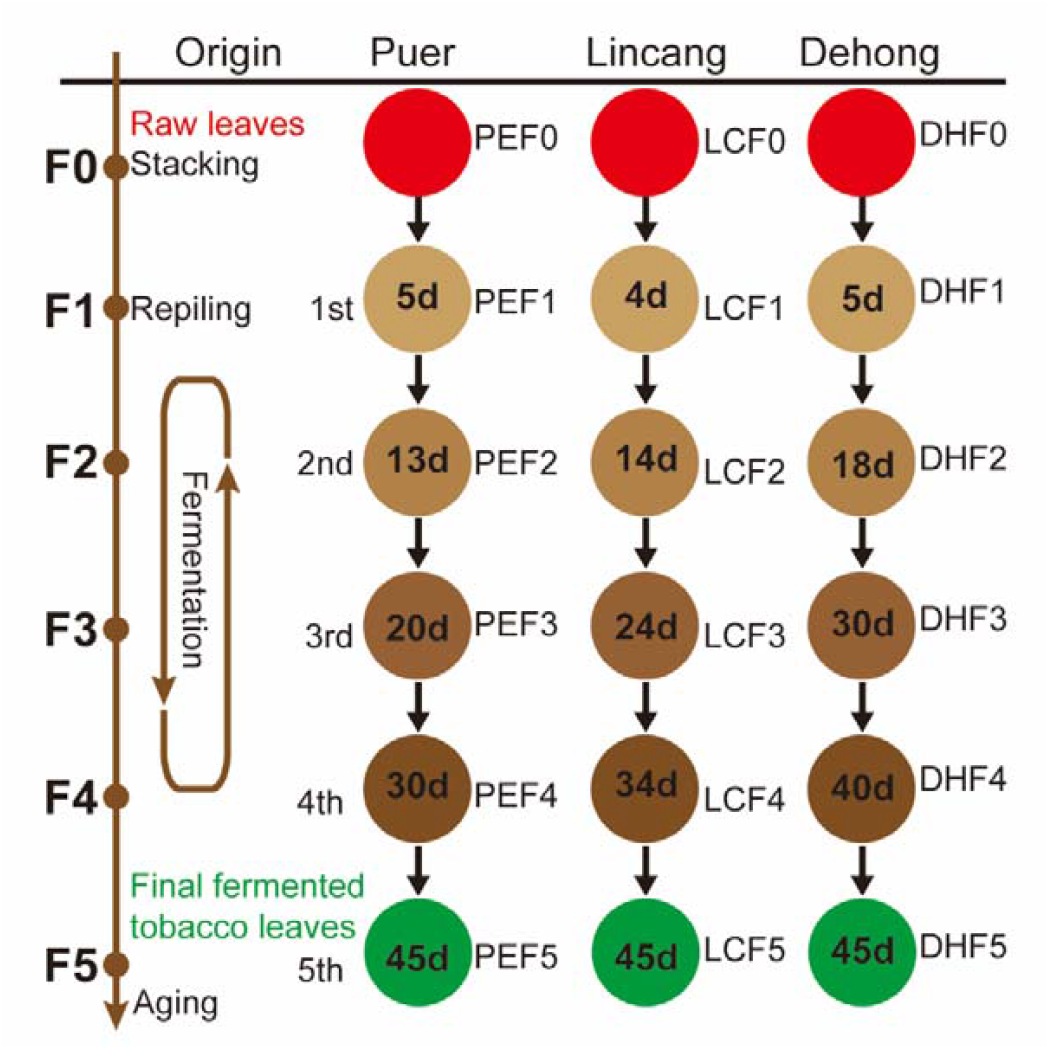
Overview of the cigar tobacco leaf fermentation processing experiments.

**FIG 2.**
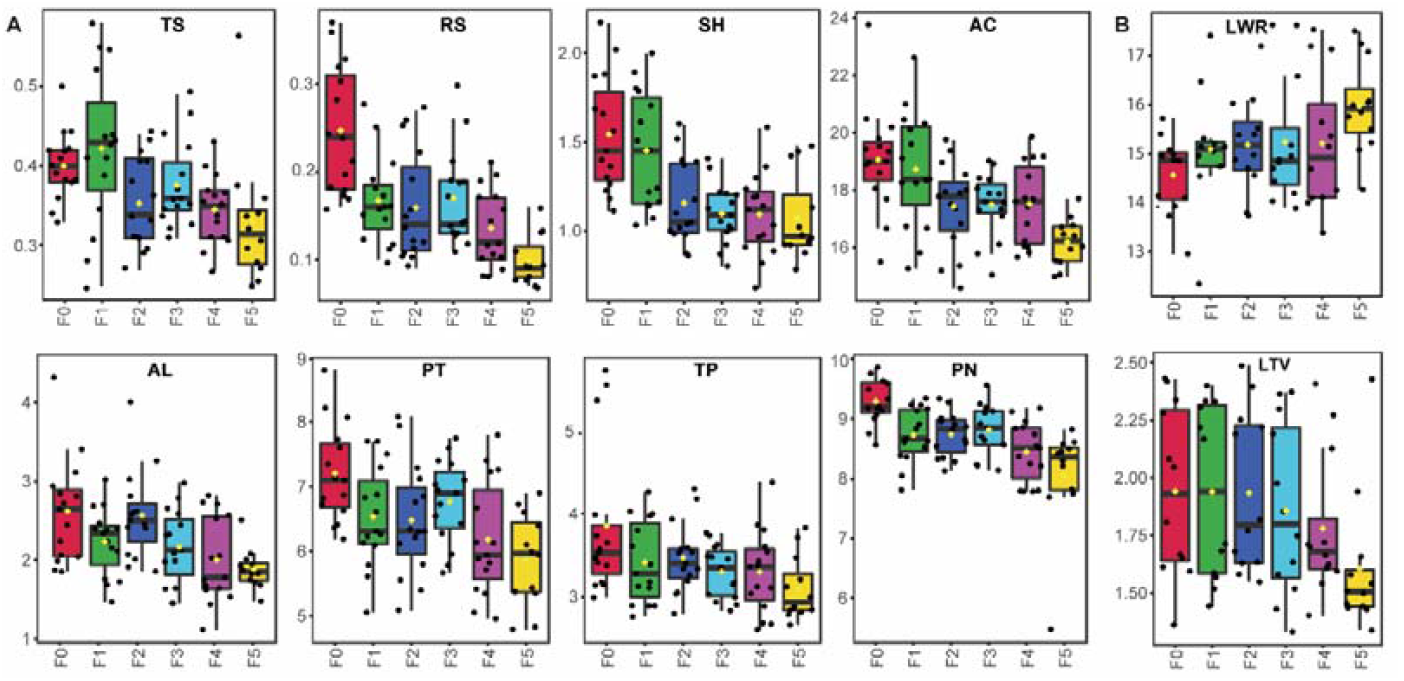
Changes in conventional chemical components(A) and physical properties(B) in cigar tobacco leaves during fermentation. TS: total sugar, RS: reducing sugar (RS), SH: starch, AC: acid cellulose, AL: acid lignin, PT: pectin, TP: total polyphenols (TP), PN: protein), LWR: leaf water retention, LTV: leaf tension. The values in the figure are the mean value of three biological replicates sourced from three locations (n=9, *P*<0.05).

### Physical properties changed during fermentation

There were no significant changes in the physical properties including leaf filling value (LFV), leaf stem ratio (LSR), leaf equilibrium moisture content (water retention, LWR), leaf thickness (LTH), leaf tension (LTV), leaf density (leaf mass weight, LMW) during fermentation (Fig. S1). However, with the prolongation of the fermentation period, the LWR showed a trend of gradual increase, and the LTV showed a trend of gradual decrease after repiling three times (Fig. 2B).

### Volative aroma compounds changed during fermentation

A total of 44 volatile aroma compounds (VACs) were further analyzed qualitatively and quantitatively based on GC–MS before and after the fermentation of cigar tobacco leaves. The VAC profiles were found to vary significantly during fermentation in Puer. Samples taken before and after fermentation could be separated obviously by PCA pattern discriminant analysis based on their VACs, with certain regularities in position distance variation (Fig. 3A). The hierarchical clustering heatmap (Fig. 3B) of VACs shows that the entire fermentation process could be divided into the early stage (F0), the middle stage (F1-F4), and the late stage (F5). In this study, orthogonal projections to latent structures discriminant analysis (OPLS-DA) analyses were performed to investigate the discriminatory VACs contributing to distinguishing the different locations before and after fermentation (Fig. 3C). A total of 19, 26 and 25 discriminatory VACs (VIP > 1, fold change > 1) were screened out from the F0 and F5 groups in DH, PE and LC, respectively (Fig. 3C). Among these, 9 volatile aroma compounds were potential markers among three locations, 3 were significantly up-regulated and 6 were significantly down-regulated, including furfural, neophytadiene, pyridine, benzyl alcohol, geranyl acetone, 2-butanone, 3-hydroxy-, N-hexanal, Isoamylalcohol and 2,3-pentanedione (Fig. 3D).

**FIG 3.**
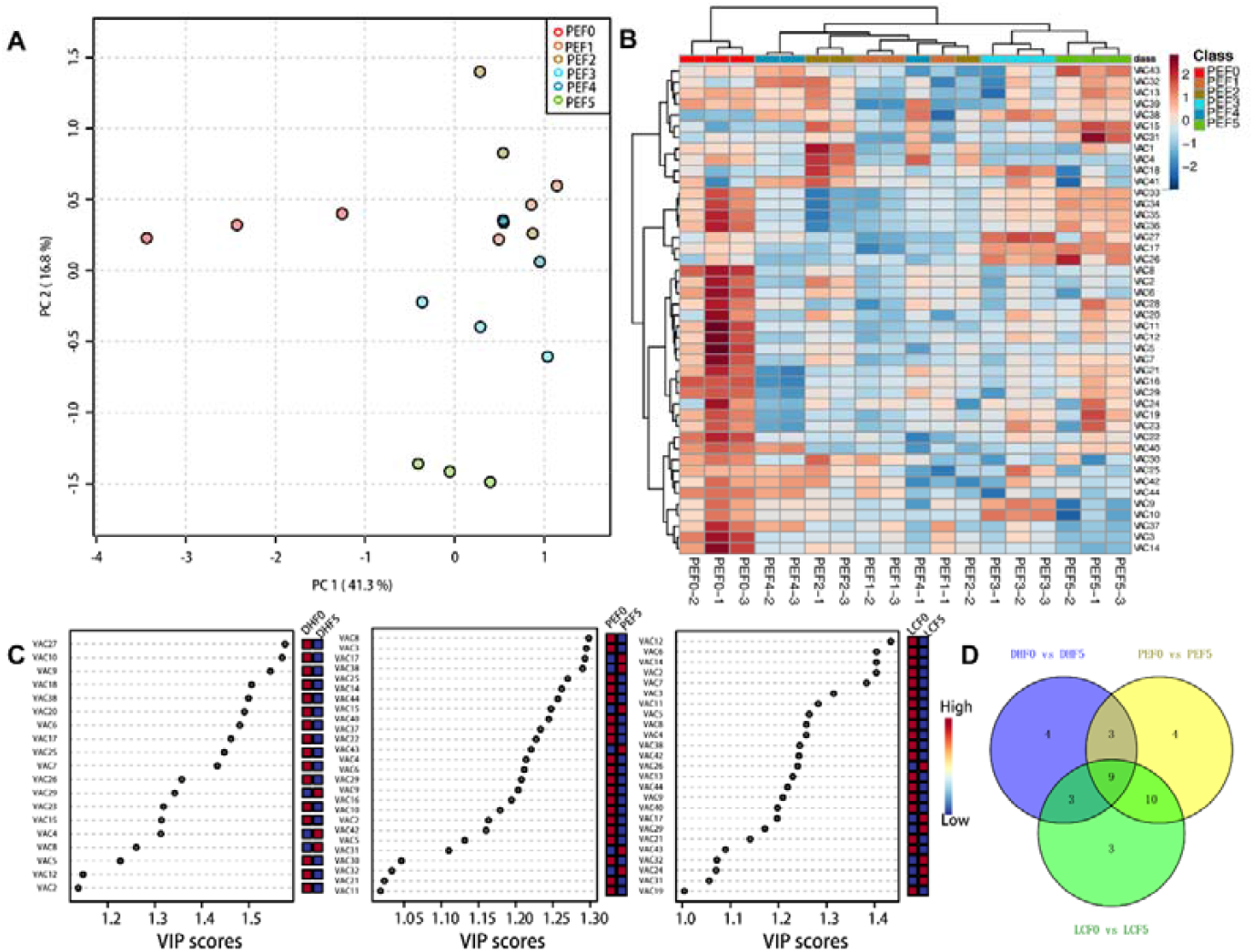
Multivariate statistical analysis of volatile aroma compounds (VACs) during fermentation of cigar tobacco leaves. (A) Score scatter plot for the principal component analysis (PCA) model (PE). (B) Hierarchical clustering analysis and heatmap visualization of VAC profiles at different fermentation stages of PE cigar tobacco leaves. (C) Important characteristics of different locations (DH, PE and LC) before and after fermentation identified by OPLS-DA. The colored boxes on the right indicate the relative concentrations of the VACs in each group under study. (D) The potential markers responsible for the aroma differences among DH, PE and LC. The screening criteria were VIP > 1 and FC > 1.

### Diversity of the microbial community in the cigar tobacco leaves

This work aimed to evaluate the changes in the microbial community of cigar tobacco leaves during fermentation stages. A total of 4,091,288 bacterial 16S rRNA and 4,197,729 fungal ITS sequences were obtained from 54 samples of cigar tobacco leaves, and the rarefaction curve tended to be flat, indicating that deep sequencing provided good overall operational taxonomic unit (OTU) coverage. Among all the OTUs identified in this study, the bacteria shared by all three locations had 111 OTUs, mainly at the genus level of *Acinetobacter, Sphingomonas, Enterobacterales, Stenotrophomonas, Bacillus, Corynebacterium, Pseudomonas* and *Staphylococcus*. PE, LC and DH had 4152, 743, 89 unique OTUs, respectively (Fig. S2A). For fungi, three locations shared 111 OTUs, mainly at the genus level of *Aspergillus, Penicillium, Alternaria, Cladosporium* and *Fusarium*. PE, LC, and DH had 337, 454 and 149 unique OTUs, respectively (Fig. S2 B).

### Microbial succession during fermentation

The diversity and similarity of the microbial community composition in the cigar tobacco leaves varied with the fermentation stages and locations. The α-diversity results showed that the diversity (Shannon index) of bacteria initially increased and then decreased. The fungi communities in our study tended to fluctuate continuously in α-diversity (Fig. 4A). The fungi α-diversity of the PE (F (5,12) =4.188, *P*=0.02) and the bacterial α-diversity (F (5,12) = 41.93, *P*< 0.001) was significantly reduced (Tab. 1). but the fungi and bacteria α-diversity of the LC and DH did not change significantly. The fungi and bacteria α-diversity did not change significantly in three locations (Tab. 1). With the extension of fermentation period, the α-diversity of fungi increased gradually and then reached their maximum value at the fourth repiling (F4) stage (F (2,6) = 36.29, *P*<0.001). The α-diversity of bacterial increased gradually and reached their maximum value at the third repiling(F3) stage (F (2,6) = 326.8, *P*<0.001).

**Tab. 1.**
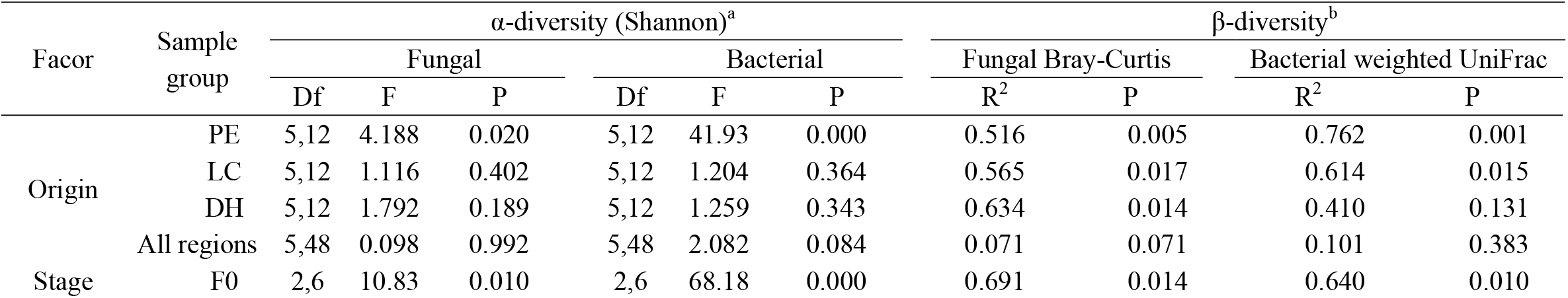
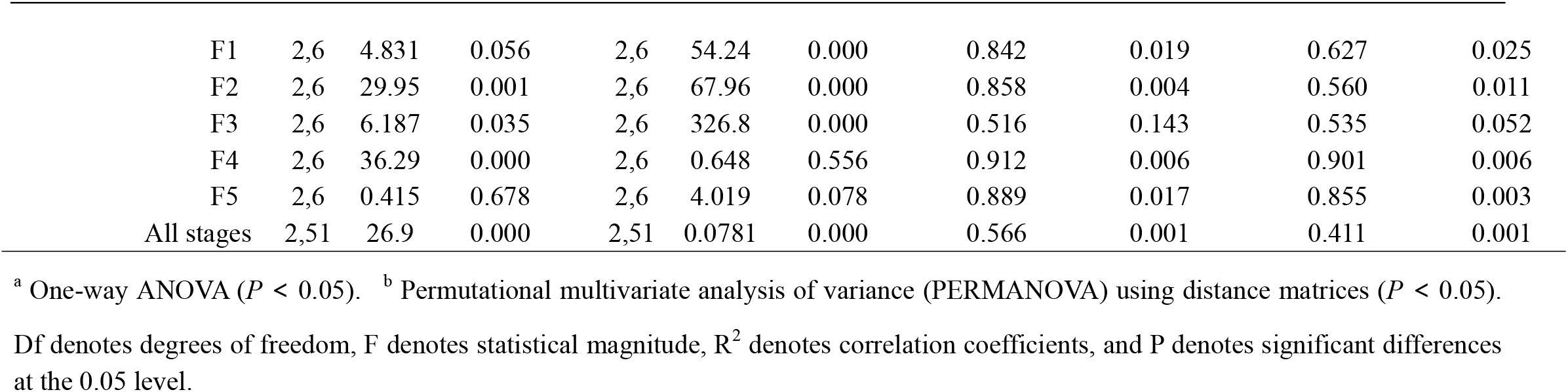
Experimental factors predicting α- and β-diversity of microbial communities in cigar tobacco leaves sourced from different locations.

**FIG 4.**
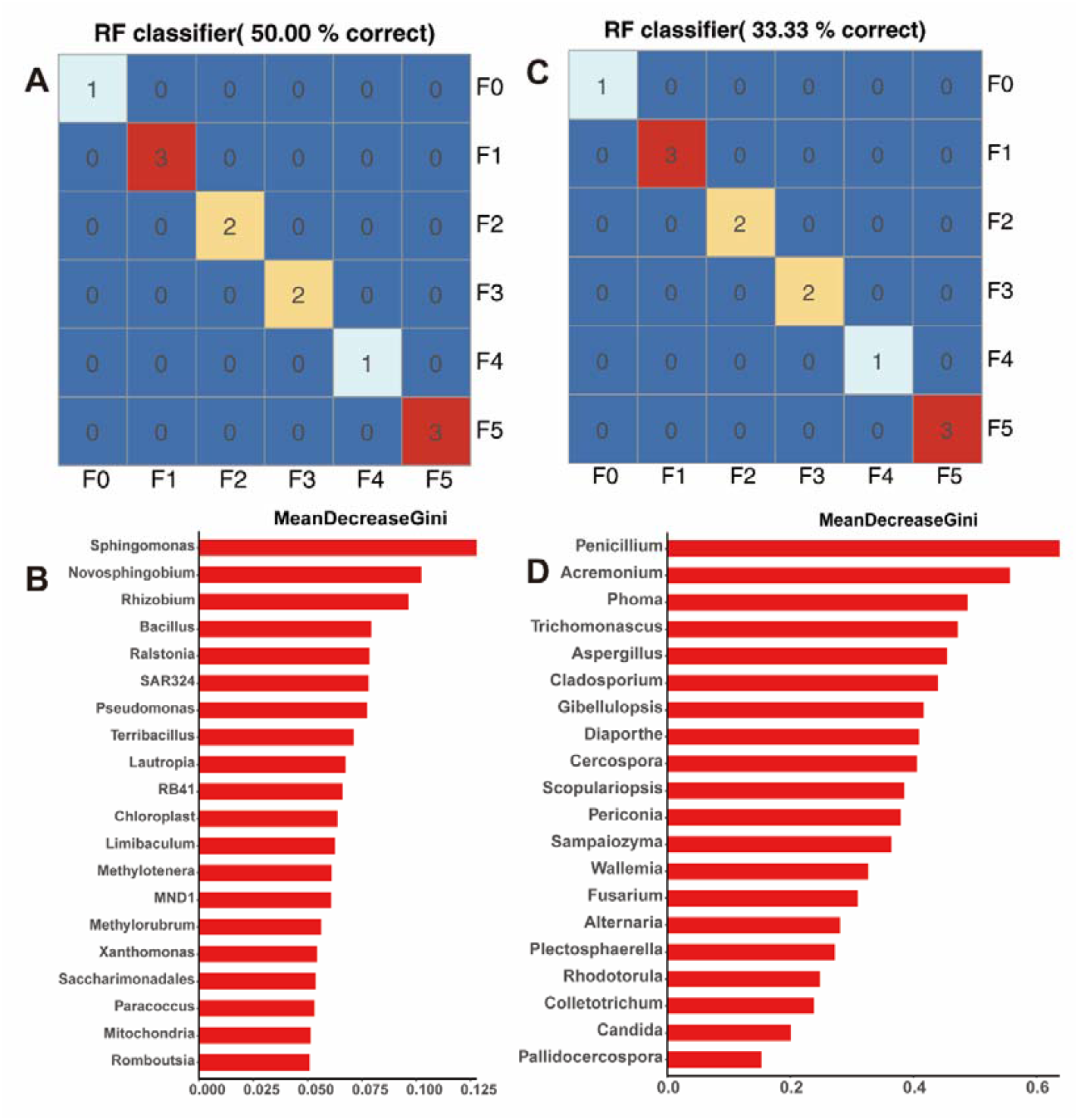
The changing trend of fungal and bacterial alpha diversity (Shannon index) during fermentation(A). The PCA graphs of the fungal community and the bacterial community at different fermentation stages and from different locations (B).

To evaluate the compositional differences of the microbial communities in the samples between locations at each stage, a principal coordinate analysis (PcoA) was performed using the weighted UniFrac metric (bacterial) and Bray-Curtis metric (fungal). Significant differences were observed in the β-diversity of fungi (R^2^ = 0.566, *P* = 0.001) and bacterial (R^2^ = 0.411, *P* = 0.001) communities at every single stage, and the locations had a little influence on the beta diversity (fungi, R^2^ = 0.071, *P* = 0.071; bacteria, R^2^ = 0.101, *P* =0.383) (Tab. 1). The microbial community structures of both bacteria and fungi had a significant temporal succession pattern between the before(F0) and after(F5) fermentation stages, but there was no obvious separation among the F1, F2, F3 and F4 fermentation stages, or in the community structure among DH, PE and LC (Fig. 4 B). The succession of the bacterial community was greater than the succession of the fungal community during fermentation (Fig. 4, Tab. 1).

### Dynamic changes in the core microbiota during fermentation

We first identified the core microorganisms of cigar tobacco leaves from three locations at different fermentation stages, which consisted of the most important taxa based on the abundance and occupancy distribution. There were significant differences in the number of core microbial communities in different producing areas, and the relative abundance and occupancy of these core members in different fermentation periods also varied (Table S2). The bacteria mainly included *Staphylococcus, Pseudomonas, Ralstonia, Sphingomonas, Bacillus, Massilia, Fibrobacter* and so on (Table S2 a, c, e). The main fungi were *Aspergillus, Cladosporium, Trichomonascus, Alternaria, Penicillium*, and *Fusarium* (Table S2 b, d, f). More bacteria than fungi were observed in these core members.

In tracking temporal dynamics and succession of core taxa during cigar tobacco leaf fermentation. The relative abundances of some core taxa had similar variation trends among PE, LC and DH, generally showing a trend of first increasing and then decreasing. However, the accumulated relative abundances were significantly different at different fermentation stages (Fig. 5). For example, the dominant bacterial genera *Pseudomonas* and *Staphylococcus* were present in high abundance in all fermentation stages of PE, LC and DH (Fig. 5A, B, C). Fig. 5D-F reveals fungal dynamics and succession during fermentation. After fermentation commenced, the relative abundance of many core fungal taxas including *Aspergillus* gradually increased, reached their maximum at the F3 and F4 fermentation stages, and then sharply decreased.

**FIG 5.**
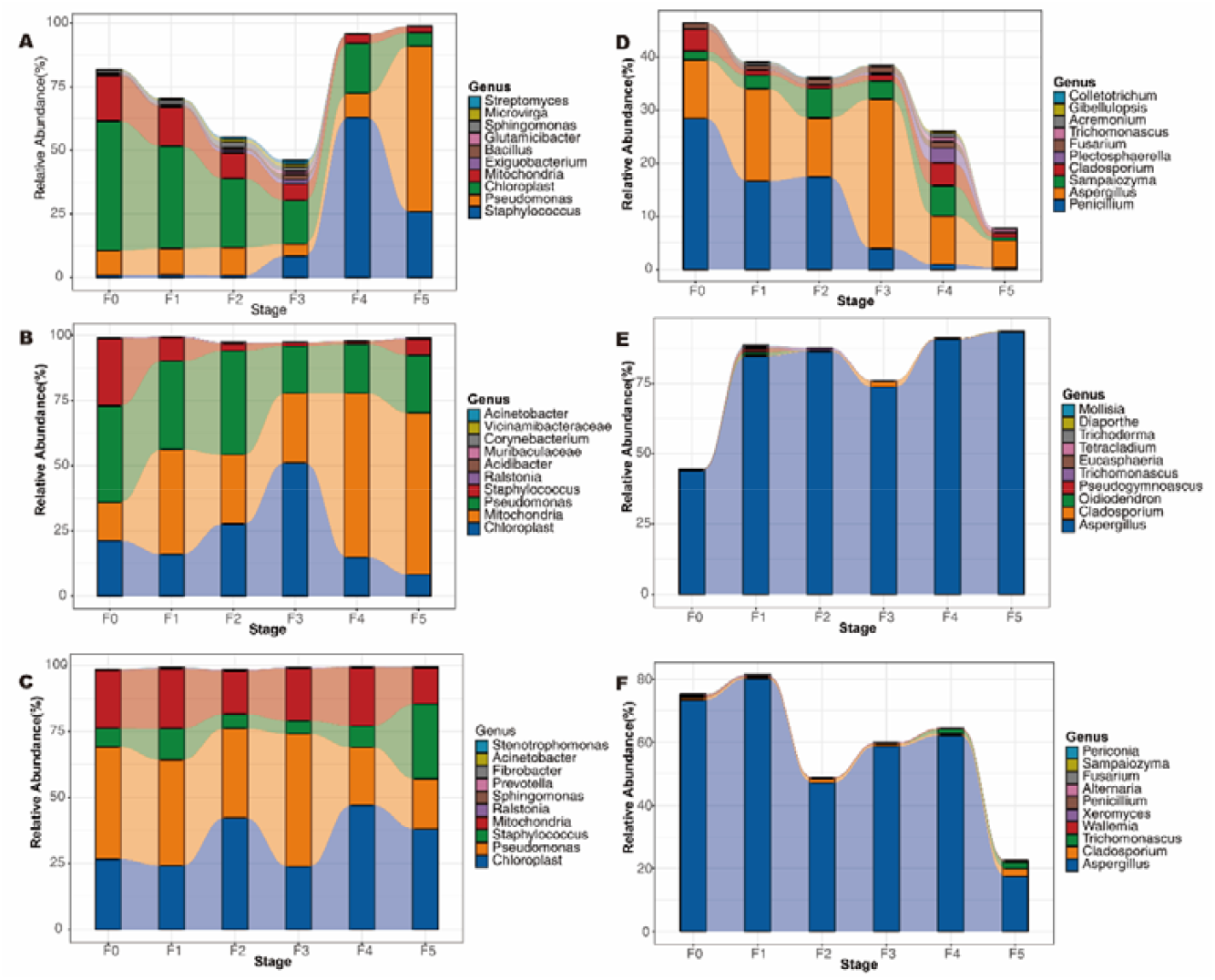
The relative abundance of core microbiota at different fermentation stages. The top 10 relative abundances are shown. A, B, and C represent the relative abundance of core bacterial communities in PE, LC, and DH, respectively. D, E, and F represent the relative abundance of core fungal communities in PE, LC, and DH, respectively.

To confirm the stability of the core taxa in different fermentation cycles and to establish a model to correlate the composition of the microbiota with the fermentation cycles, the random forest supervised learning model was used to regress the relative abundance of bacteria and fungi at the genus level against the fermentation time. The model results showed that bacterial core taxa had better discrimination than fungi and could correctly identify all samples of the PE. The model explained 50.00% of the microbiota variance associated with the fermentation process of PE (Fig. 6A, C). Discrimination of the LC and DH was low, reaching only 33.33% and 16.67%, respectively (Fig. S3A, S4A). The fungal core taxa had lower resolutions for the PE, LC and DH samples, reaching 33.33%, 25.00% and 16.67%, respectively (Fig. 6C, Fig. S3C, Fig. S4C). The Gini index model showed that many core taxa were important features of the fermentation period model. For example, *Sphingomonas* and *Penicillium* could explain the largest variation of bacterial and fungal communities in PE locations, respectively (Fig. 6B, D). *Cupriavidus* and *Trichomonascus* could explain the largest variation in bacterial and fungal communities in LC locations, respectively (Fig. S3B, D). *Corynebacterium* and *Aspergillus* could explain the largest variation in bacterial and fungal communities in DH locations, respectively (Fig. S4B, D). Taken together, these results confirmed that these microorganisms might be biomarkers associated with certain stages of fermentation and may reflect the functional characteristics at different stages.

**FIG 6.**
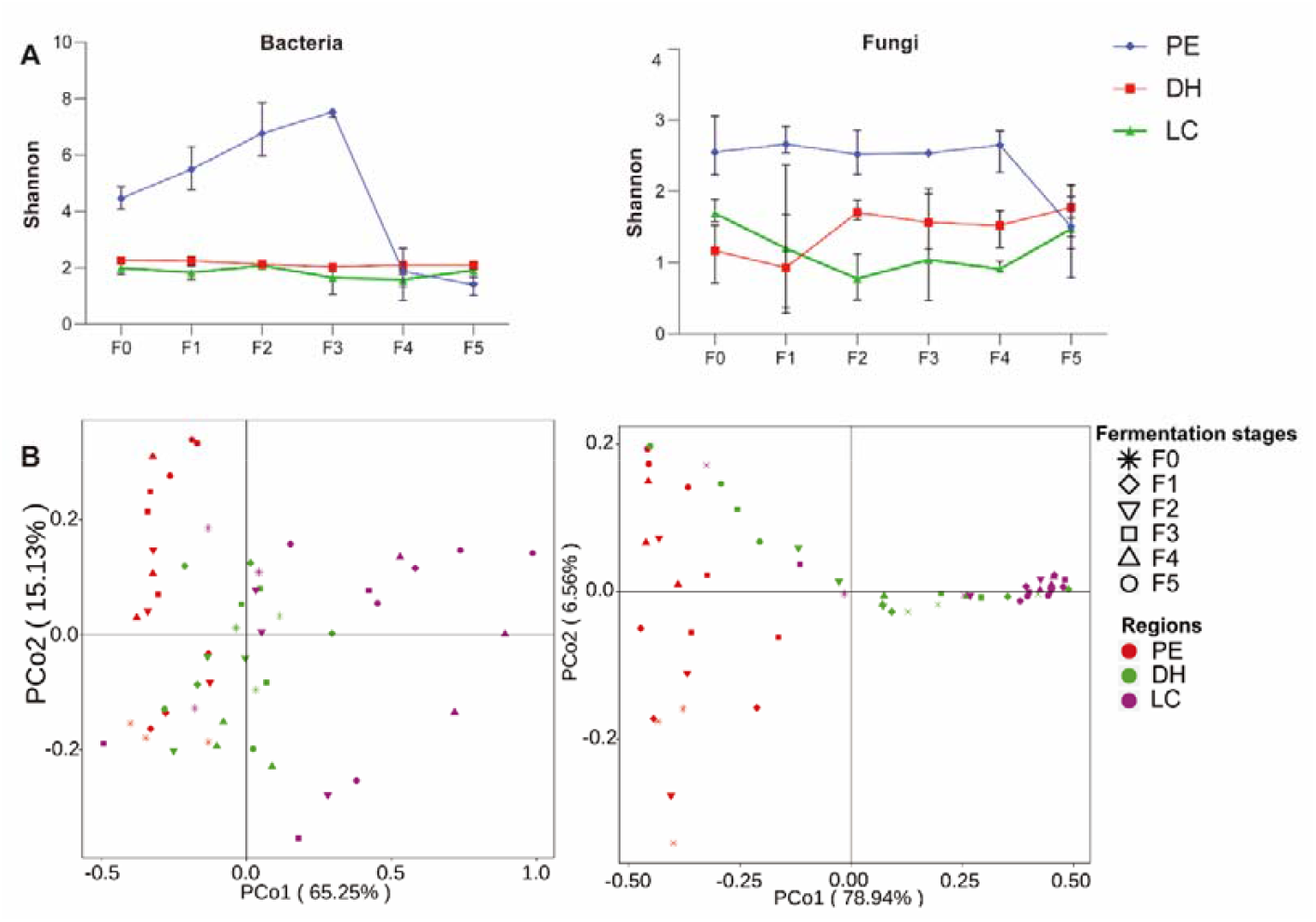
The core taxa can distinguish the fermentation stages of cigar tobacco leaves. Classification of random forest models of the fermentation stage of the core taxa of PE bacteria (A) and fungi (B). The important features (top 20) based on Mean Decrease Gini (MDG) of random forest models of the core taxa of PE bacteria (C) and fungi (D).

### Relationship between physicochemical characteristics and microorganisms during fermentation

To investigate the relationship between metabolic enzymes, physicochemical properties and microbial community composition. Canonical correspondence analysis (CCA) was performed based on microbial abundance, metabolic enzymes and physicochemical properties. The results showed that there was a coupling between the microbial communities, enzymes and metabolic physicochemical properties. The measured metabolic enzymes and physicochemical properties could explain 35.25% and 42.77% of the bacterial community variation, respectively (Fig. S5 A, B). The measured metabolic enzymes, physicochemical properties and volatile components could explain 45.88% and 42.00% of the fungal community variation, respectively (Fig.S5 D, E). The metabolic enzymes: AL, NAD-MDH, PAL, LiP and physicochemical properties pH, PEE, TAA, PN, NT, Mg, K were the most important factors affecting the bacterial community, with AL, pH, PEE, TAA and PN being the most important factors. The metabolic enzymes AL, NAD-MDH, GDH, PAL, LiP, NR and CAT, and the physicochemical properties pH, PEE, TAA, NT, TS, TN, Mg were the most important factors affecting the fungal community, with AL, NAD-MDH, NR, pH, NT, TS, TN, and Mg being the most important factors (Fig. S5). Overall, multiple metabolic enzymes and physicochemical properties were likely to have a collective effect on the microbial community.

Furthermore, we studied the bacterial and fungal community functions in the samples and determined the different functions of the communities in the different groups. The results showed that the main functions of core bacteria were carbohydrate metabolism, glycan biosynthesis and metabolism, energy metabolism, amino acid metabolism, etc. The functions of the core fungi are mainly saprotrophs, pathotrophs and symbiotrophs (Fig. 7A). To further explore the co-occurrence patterns between core taxa and metabolic enzymes, physicochemical properties and volatile compounds, molecular ecological networks were constructed. In general, a more densely connected module was observed in the bacteria than in the fungi (Fig. 8, Fig. S6). The network of bacteria and volatile aroma components, physicochemical properties, and metabolic enzymes consisted of 181, 38, and 21 associations, respectively, in which 113 edges were positive associations. Most core taxa belong to *Aureimonas, Ralstonia, Skermanella, Methylobacterium-Methylorubrum, Ensifer, Steroidobacter, Staphylococcus, Bacillus* and *Microvirga*. In this network, some microbial taxa had significant correlations with various volatile aroma compounds, especially cibai trienediol (VAC41-43), 1-penten-3-one (VAC1), 2-butanone, and 3-hydroxy (VAC4), which had high connectivity (Fig. 8 A). Some microbial taxa had high connectivity with acid lignin (AL), nicotine (NT), amylase (AL), glutamate dehydrogenase (GDH), and malic dehydrogenase (NAD-MDH) (Fig. 8C-F). The network of fungi and volatile aroma compounds consisted of 99 associations (Fig. 8 B), 69 edges of which were positive associations. Most core taxa belonged to *Aspergillus, Penicillium, Sampaiozyma, Trichomonascus, Fusarium, Wallemia, Cercospora, Pallidocercospora, Mortierella, Xeromyces, Filobasidium, Vishniacozyma, Sporobolomyces* and *Podospora*. In general, some bacteria and fungi jointly participated in the metabolism of the fermentation process and played different roles.

**FIG 7.**
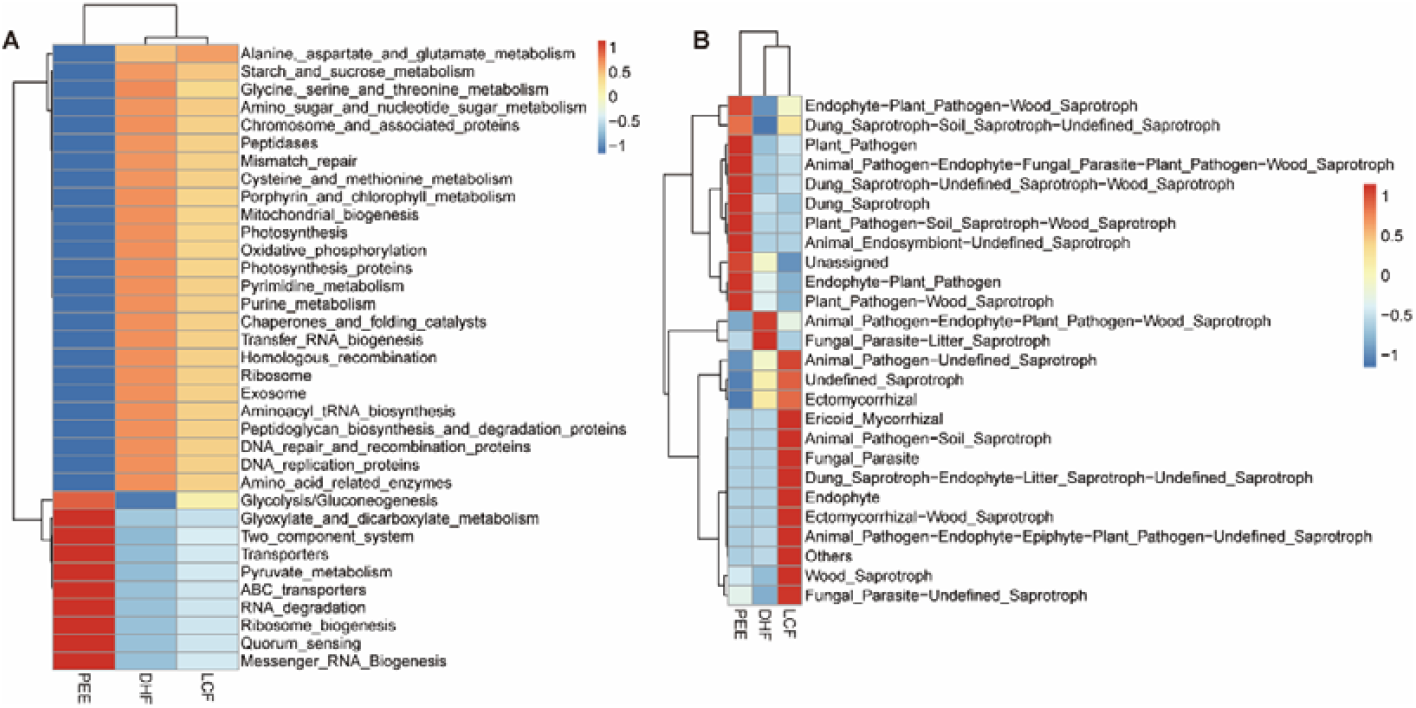
Variations in bacterial (A) and fungal (B) functional profiles during fermentation were analyzed using Tax4fun and FUNGuild.

**FIG 8.**
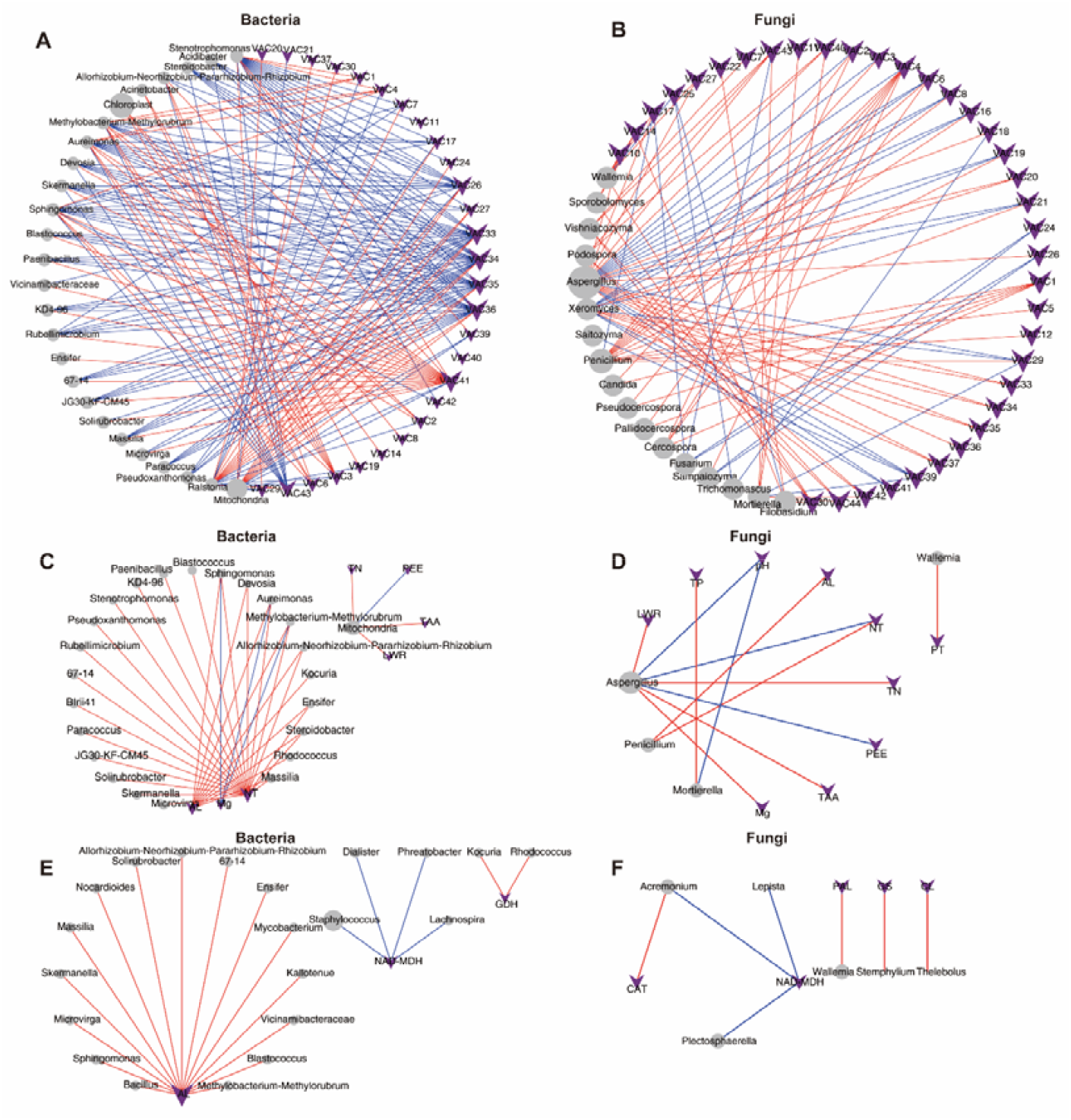
Network analysis based on the cooccurrence of volatile components (A, B), physicochemical properties (C, D), metabolic enzymes (E, F) and the bacterial and fungal communities. Purple vee nodes represent volatile components. Grey ellipse nodes represent microbial members. Direct connections between nodes indicate strong correlations (Pearson correlation coefficient, *P* < 0.05). Red lines represent positive interactions between nodes, and blue lines represent negative interactions. The sizes of vee nodes represent the interconnected degree. The sizes of the circle nodes represent the average relative abundances of bacteria and fungi.

## DISCUSSION

Fermentation is considered the process of biomass degradation and transformation, which affects the nutritional content, bioactivity, aroma, and flavor properties of the product (15).Various metabolic pathways were involved in the fermentation process of cigar tobacco leaves, including biosynthesis of nonnitrogen compounds, degradation and transformation of nitrogen compounds, and the Maillard reaction. For example, organic matter in tobacco leaves will produce inorganic products such as carbon dioxide, water, and ammonia, by combining hydrogen and oxygen in the process of dehydrogenation or degradation. Secondary alkaloids, organic acids, aromatic oils and other volatile substances were released and discharged, resulting in the reduction of dry matter (2). In this study, the physicochemical properties and volatile aroma compounds of cigar leaves from three locations were analyzed. The results showed that nonnitrogen compounds including total sugars, reducing sugars, starch, pectin, total polyphenols, lignin and cellulose decreased by 11.37%, 63.02%, 30.58%, 15.84%, 20.95%, 32.91% and 17.05%, respectively. Nitrogen compounds, such as protein, total nitrogen and nicotine decreased by 11.73%, 2.94% and 4.21%, respectively, while amino acids increased by 18.77% (Fig. 2A, Fig. S1). Similar results were demonstrated in a recent study in which dominant bacteria and fungi in tobacco leaves had important application prospects in the degradation of carbohydrates, proteins, cellulose, lignin, aromatic compounds and aliphatic compounds (2, 4). Therefore, it is important to explore the key microorganisms in the process of biomass fermentation, find an efficient enzyme system, and clarify the molecular mechanism of microbial degradation and their interaction relationship.

At the initial stage of fermentation, the contents of TS, RS and SH decreased sharply due to the metabolism of biodegradable organic compounds by bacteria and fungi, producing CO_2_, H_2_O and organic acids. Pectin is decomposed under the catalysis of pectinase, converted into methanol and volatilized (2). Previous studies have found increased lignin methoxyl contents in tobacco leaves after fermentation. and a portion of cellulose was oxidized, contributing to the changes in tension of fermented tobacco leaves (1), Our results also showed that leaf tension increased at the beginning and then decreased during fermentation, which was consistent with previous studies (Fig. 2B). Tao et al.(13) found that *Pseudomonas putida* and *Sphingomonas* in tobacco leaves could produce hydrolytic enzymes to degrade cellulose through their own metabolism. In our study, the dominant genera in tobacco leaves were distributed in *Pseudomonas* and *Sphingomonas* (Fig. 5). Consistent with our study (Fig. 2), Frankenbur (2) found that the tannin and polyphenol contents of cigar leaves decreased from 0.48% to 0.36% after fermentation, and the polyphenols were oxidized to form quinones, which can interact with amino acids and form melanin substances. The changes in pH values are shown in Fig. S1. The initial pH value was 6.17, which then increased to 6.53 after fermentation. The increase in the pH value may be due to the release of large amounts of NH_4_^+^-N by bacteria during the fermentation process in the decomposition organic matter, such as proteins and nitrogen-containing bases(16). Consistently, nitrogen-containing compounds were volatilized or degraded and then transformed from water-soluble nitrogen compounds to insoluble nitrogen during fermentation. The genus *Bacillus* is widely distributed in the natural environment and members of *Bacillus* provided about 35% of the microbial protease enzymes in the world. Proteases such as collagenase have recently been discovered and isolated from *Bacillus* cereus (13). In particular, macromolecular proteins in tobacco leaves are decomposed into amino acids, amides and quinones. Soluble nitrogen-containing substances, including amine nitrogen and amide nitrogen, decreased sharply, and a large amount of ammonia was produced after oxidative deamination, which was gradually volatilized out of tobacco leaf tissues during fermentation. The remaining amino acids reacted with quinone substances in tobacco leaves to produce macromolecular polymers. Nicotine is oxidized by fermentation to produce secondary alkaloids and pyridine derivatives (17-20). Consistently, our results showed that the main core bacteria *Bacillus* maintained high abundance at different fermentation stages (Fig. 5), and CCA and cooccurrence network results showed that *Bacillus* was significantly correlated with physicochemical properties (Fig. 8, Fig. S3, 4, 5). Tobacco leaves are full of colloidal capillary porous substances, and colloidal substances (such as protein, pectin, cellulose, etc.) and crystalline substances in the contained substances determine the hygroscopicity of tobacco leaves. After fermentation, some osmotic crystalline substances (such as water-soluble sugars, organic acids, inorganic salts, etc.) and macromolecular colloidal substances were degraded, resulting in a decrease in the self-heating and hygroscopic effects of tobacco leaves(14), which explains the results of this study, that is, with the extension of the fermentation period, the equilibrium moisture contents of tobacco leaves gradually increased.

Tobacco leaves are rich in aromatic amino acids and terpene compounds, which are important aroma precursors. Their degradation products are one of the important aroma sources in tobacco(21). After fermentation, the total amount of VACs were increased (8). In this study, we observed 16 kinds of important features in different fermentation stages, including valeraldehyde, 6-methyl-2-heptanone, furfural, solavetivone, cibai trienediol-4, beta-ionone 5,6-epoxide, N-hexanal, pyridine, cyclohexanone, 2,3-dimethyl-2-(3-oxobutyl)-, isoamylalcohol, benzaldehyde, 5-hepten-2-one, 6-methyl-, 2,3-pentanedione, 2-acetyl-1-pyrroline, 2-furanmethanol, 2,6-nonadiendienal, (E,E), cibai trienediol-3, consisting of seven nonenzymatic browning reaction products, three cibai alkane degradation products, four carotenoid degradation products, one alkaloid and one heterocyclic. Furthermore, there were significant differences in the volatile aroma compounds before and after fermentation in different locations (Fig. 3). The microbial community was closely related to VACs, and aldehydes and ketones were positively related to *Staphylococcus, Corynebacterium, Tetragenococcus, Yaniella*, and *Enteractinococcus* (22). Our results also showed that VAC33, VAC34 VAC35, VAC36 and VAC41 were significantly associated with a variety of bacteria, and the connection between fungi and VACs was less than the connection of bacteria (Fig. 8), indicating that the cigar tobacco leaves produced under different ecological conditions have different microbial communities also lead to the difference in VACs and aroma characteristics. In our study, Tax4Fun and FunGuild were used to evaluate the function of bacterial and fungal communities. In the process of cigar tobacco leaves fermentation, bacterial communities were found to be annotated to a variety of functions, including carbohydrate metabolism, amino acid metabolism, lipid metabolism, and glycan biosynthesis and metabolism. Fungi are mainly annotated to saprotroph action. The results show that a variety of bacteria can produce highly active enzyme systems in the growth and reproduction process. Under the synergistic action of enzymes, chemistry and microorganisms, the macromolecular substances in tobacco leaves can be degraded, oxidized, reduced, polymerized, coupled and transformed to form low molecular weight compounds, mainly including various aroma compounds. The saprophytic action of fungi degrades lignin, cellulose and pectin in tobacco leaves to form small molecules such as glyoxal and glucose.

Microbial degradation and transformation of organic compounds play a crucial role in the fermentation process(13). Microorganisms (including bacteria and fungi) are the main participants in the conversion of materials during the cigar tobacco leaf fermentation process(4). Our data showed that there were significant differences in the species and structure of microbial communities on cigar tobacco leaves sourced from different locations in Yunnan. There were more bacterial species than fungi, and PE had the most abundant bacterial communities, while DH had the limited fungal and bacterial species (Fig. S2). Consistent with our findings, previous studies have shown that the main microbial communities of tobacco leaf fermentation include *Pseudomonas, Bacillus, Methylobacterium, Acinetobacter, Sphingomonas, Neophaeosphaeria* and *Cladosporium* (12). Microorganism play an important role in the fermentation process of cigar tobacco leaves, and are affected by moisture, temperature, enzymes and physicochemical properties. The CCA and cooccurrence network results showed that some microbial taxa had significant correlations with various metabolic enzymes and physicochemical properties (Fig. 8, Fig. S5, Fig.S6). As bacteria or fungi produce various active enzymes in their metabolism, these enzymes are secreted extracellularly and degrade macromolecular organic compounds in tobacco leaves (9, 23, 24). Various strains of the *Bacillus* genus can secrete cellulase, pectinase, protease and other functions(25), improving the aroma quality of cigar tobacco leaf fermentation and accelerating sweating.

Cell wall substances (such as cellulose and pectin) and nitrogen-containing compounds in tobacco leaves have a great influence on the physical properties and intrinsic quality of tobacco leaves. This study found that nonnitrogen compounds including total sugars, reducing sugars, starch, pectins, total polyphenols, lignins and cellulose were significantly decreased during fermentation. The macromolecular nitrogen compounds such as protein and nicotine decreased significantly, while micromolecular nitrogen compounds such as amino acids increased significantly. The furfural, neophytadiene, pyridine, benzyl alcohol, geranyl acetone, 2-butanone, 3-hydroxy, n-hexanal, isoamylalcohol and 2,3-pentanedione were potential markers affecting the aroma quality before and after fermentation. The microbial diversity of the cigar tobacco leaves in PE, LC and DH of China were initially increased and then decreased during fermentation. This diversity was mainly associated with fermentation time. At the same time, the microbial diversity of different locations was significantly different. These findings indicated that phylloplane microorganisms play a vital role in the quality and regional characteristics of cigar tobacco leaves.

## MATERIALS AND METHODS

### Experimental materials

Cigar tobacco (*Nicotiana tabacum*, Yunxue No. 1) was grown in Lincang (LC), Puer (PE) and Dehong (DH) in Yunnan. The freshly harvested cigar tobacco leaves were strung on a bamboo or wooden pole and air-curied in a local curing room for 40 days. After air-curing in DH and LC, the tobacco leaves were naturally fermented locally for one month. The tobacco leaves in the PE were immediately transported to the fermentation factory after curing. Central tobacco leaves of 1000 kg were selected from three locations and sent to the fermentation factory in Yuxi for standardized artificially controlled fermentation. The fermentation experiments were carried out according to Yunnan cigar tobacco leaf stacking fermentation technical regulations (enterprise standard, Q/YNYC(KJ). J02-2022), and the specific treatment process includes pest control, sorting, rehumidification, balancing, stacking, fermenting, repiling (5 times) and unstacking. Turning occurred immediately when the temperature reached 40°C or when there was a cooling trend. Approximately 500 g of tobacco leaves taken before fermentation(F0) and during each repiling (F1, F2, F3, F4, F5) were used in the following analyses. Each sample was given a specific code as shown in Fig. 1. The growing locations, including Lincang, Puer and Dehong, were denoted LC, PE and DH, respectively. Fermentation samples were denoted followed by a number (indicating the fermentation stages or the numbers of repilings). A total of 54 cigar tobacco leaf samples from three origins and six different fermentation cycles were collected. Approximately 20 g of cigar tobacco leaves were cut with sterilized scissors, placed in centrifugal tubes quickly frozen and stored at -80°C for DNA extraction and enzymatic analysis. The remaining tobacco leaves were used for chemical composition analysis, physical property evaluation, and aroma compound determination. Three independent biological replicates were analyzed for each location or variety.

### Determination of chemical compositions

The indices of the main chemical compositions were total sugar (TS), reducing sugar (RS), total nitrogen (TN), protein (PN), starch (SH), nicotine (NT), potassium oxide (K), chloride ion (Cl), magnesium (Mg), petroleum ether extract (26), pectin (PT), acid cellulose (AC), acid lignin (AL), total polyphenols (TP), total amino acids (TAA) and pH. The content of each index was determined by using a continuous flow analytical system or ion chromatography (27). The contents of TS, RS, TN, PN, SH, NT, K, Cl, Mg, PEE, PT, AC, AL and pH were separately determined in accordance with the standards in the tobacco industry YC/T 159-2019, YC/T 159-2019, YC/T 161-2002, YC/T 249-2008, YC/T 249-2008, YC/T 468-2013, YC/T 217-2007, YC/T 162-2011, YC/T 175-2003, YC/T176-2003, YC/T 346-2010, YC/T 347-2010, YC/T 347-2010, YC/T 222-2007. The contents of TP and TAA were determined using a spectrophotometric method based on the books published by Yu (2009).

### Determination of physical properties

Physical indicators of cigar tobacco include leaf filling value (LFV, cm3·g-1), leaf stem ratio (LSR, %), leaf water retention (LWR, %), leaf thickness (LTH, μm), leaf tension (LTV, N), leaf density (leaf mass weight, LMW, %). All the physical index methodologies were based on the books published by Yu (28).

### Determination of volatile aroma compounds

The samples of LCF0, LCF5, DHF0, DHF5, PEF0, PEF1, PEF2, PEF3, PEF4, PEF5 were used for the determination of aroma substances. The indices of 44 volatile aroma compounds (VACs, Table S1) were detected by steam distillation extraction and combined gas chromatography/mass spectrometry (GC-MS) (27, 29). Quantitation based on the quantitation ion which is unique in the coeluted components increased the accuracy of peak integration.

### Determination of metabolic enzymatic activity

The activities of 18 metabolic enzymes during fermentation in different producing areas were determined and analyzed. Including peroxidase, neutral protease (NP), lignin peroxidase (LiP), cellulase (CL), phenylalanine ammonia-lyase (PAL), pectinase (PN), catalase (CAT), sucrase (SU), polyphenol oxidase (PPO), total amylase (AL), α-amylase (α-AL), β-amylase(β-AL), neutral invertase (NI), AD-malate dehydrogenase (NAD-MDH), glutamine synthetase (GS), glutamate dehydrogenase (GDH), lipoxygenase (LOX), nitrate reductase (NR), and alkaline protease (AKP). All enzyme activities were determined using an enzyme-linked immunosorbent assay (ELISA) kit (Solarbio or mlbio, Beijin, China) following the manufacturer’s instructions.

### Microbial DNA extraction and sequencing

Total genomic DNA from the samples was extracted using the cetyltrimethylammonium bromide (CTAB) method. DNA concentration and purity was monitored on 1% agarose gels. The V4 regions of the bacterial 16S rRNA gene were amplified using the forward primer 515F (5 ′ -GTGCCAGCMGCCGCGGTAA-3 ′) and the reverse primer 806R (5 ′-GGACTACHVGGGTWTCTAAT-3′). The fungal internal transcribed spacer gene was amplified using the universal primers ITS1F (5 ′-CTTGGTCATTTAGAGGAAGTAA-3 ′) and ITS2R (5 ′-GCTGCGTTCTTCATCGATGC3 ′). The PCR products quantification and qualification was carried out based on previous methods (30). Sequencing libraries were generated using the TruSeq® DNA PCR-Free Sample Preparation Kit (Illumina, San Diego, CA, USA) following the manufacturer’s recommendations and index codes were added. The library quality was assessed on the Qubit@ 2.0 Fluorometer (Thermo Scientific, Waltham, MA, USA) and Agilent Bioanalyzer 2100 system (Agilent, Santa Clara, CA, USA). The library was sequenced on an Illumina NovaSeq platform and 250 bp paired-end reads were generated.

### Processing of sequencing data

Paired-end reads were split based on their unique barcodes and were cut off the barcodes and primer sequences. Using FLASH (Version 1.2.11) (31) was used to merge paired-end reads when at least ten bases overlaped with the opposite end reads of the same DNA fragment. Quality filtering were performed using the fastp (Version 0.20.0) software to obtain high-quality clean tags. The clean tags were compared with the reference database (Silva database https://www.arb-silva.de/ for 16S, Unite database https://unite.ut.ee/ for ITS) using Vsearch (Version 2.15.0) (30)to detect and remove the chimera sequences. Denoise was performed with the DADA2 or deblur module in the QIIME2 software (Version QIIME2-202006) to obtain initial ASVs (amplicon sequence variants), and then ASVs with abundances less than 5 were filtered out(32). Species annotation and multiple sequence alignment were performed using QIIME2 software. The absolute abundance of ASVs was normalized using a standard sequence number corresponding to the sample with the least sequences. Subsequent analysis of alpha diversity and beta diversity were all performed based on the output normalized data. Furthermore, Tax4Fun (33) and FUNGuild (34) were used for functional annotation analysis of bacteria and fungi, respectively.

### Data availability

The raw sequencing data have been uploaded to the National Center for Biotechnology Information (NCBI) Sequence Read Archive (SRA) database under BioProject number PRJNA856456.

### Statistical analyses

The statistical analysis was performed using the SPSS software package (Chicago, IL, USA) and MetaboAnalyst platform (https://www.metaboanalyst.ca/). Tukey’s Honestly Signicant Difference (Tukey’s HSD) was used to determine the mean differences between different fermentation stages. Student’s *t* test was used to determine if the physicochemical properties indices before fermentation were significantly different from those after fermentation. A probability of *P*< 0.05 indicated that the measured differences were significant. Alpha diversity and beta diversity estimates were calculated with QIIME2 using weighted UniFrac distance between samples for bacterial 16S rRNA reads and Bray-Curtis dissimilarity for fungal ITS reads. Principal coordinate analysis (PCoA) was performed to evaluate the distribution patterns of microbiomes based on β-diversity calculated by the Bray-Curtis distance with the LabDSV R package. Oneway analysis of variance (ANOVA) was used to determine whether fermentation stages and regions contained statistically significant differences in diversity. Permutational multivariate analyses of variance (PERMANOVA) using distance matrices to determine the statistically significant differences with the adonis function in the vegan R package. Canonical correspondence analysis (CCA) visualization diagram was drawn using an RStudio package (version 2.15.3).

## SUPPLEMENTARY MATERIAL

Figure S1. The chemical and physical important features selected by ANOVA plot (*P*<0.05).

Figure S2. This Venn diagram shows specific OTUs and shared OTUs of bacteria and fungi among different locations.

Figure S3. The core taxa can distinguish the fermentation stages of cigar tobacco leaves in LC.

Figure S4. The core taxa can distinguish the fermentation stages of cigar tobacco leaves in DH.

Figure S5. CCA reflected the relationship between metabolic enzymes, physicochemical properties, volatile aroma compounds and microbial community composition during cigar tobacco leaf fermentation.

Figure S6. Network analysis based on the cooccurrence of physicochemical properties, metabolic enzymes and the bacterial and fungal communities.

Table S1. The corresponding information of volatile aroma components determined by GC–MS analysis.

Table S2. Genus identified as core taxa communities at different fermentation stages in different locations.

## ACKNOWLEDGMENTS

This work was supported by the China Tobacco Monopoly Bureau Grants and Yunnan Provincial Tobacco Monopoly Bureau Grants (110202101012(XJ-04)/2021530000241002), and Yunnan Provincial Tobacco Monopoly Bureau Grants (2020530000241001).

## REFERENCES

1. Frankenburg WG. 1946. Chemical changes in the harvested tobacco leaf: Part I. chemical and enzymic conversions during the curing process, p 309–387, Adv Enzymol Relat Areas Mol Biol doi:10.1002/9780470122518.ch8.

2. Frankenburg WG. 1950. Chemical changes in the harvested tobacco leaf. II. Chemical and enzymic conversions during fermentation and aging. Adv Enzymol Relat Areas Mol Biol 10:325–441. doi:10.1002/9780470122556.ch8.

3. Banozic M, Jokic S, Ackar D, Blazic M, Subaric D. 2020. Carbohydrates-key players in tobacco aroma formation and quality determination. Molecules 25. doi:10.3390/molecules25071734.

4. Liu F, Wu Z, Zhang X, Xi G, Zhao Z, Lai M, Zhao M. 2021. Microbial community and metabolic function analysis of cigar tobacco leaves during fermentation. MicrobiologyOpen 10. doi:10.1002/mbo3.1171.

5. Yang Y, Peng Q, Ou M, Wu Y, Fang J. 2018. Research progress in tobacco fermentation. Journal of Biosciences and Medicines 06:105–114. doi:10.4236/jbm.2018.66008.

6. Zelitch I, Zucker M. 1958. Changes in oxidative enzyme activity during the curing of connecticut shade tobacco. Plant physiology 33:151–5. doi:10.1104/pp.33.2.151.

7. Jensen C, Parmele H. 1950. Fermentation of cigar-type tobacco. Industrial & Engineering Chemistry 42:519–521. doi:10.1021/ie50483a032.

8. Giacomo M, Paolino M, Silvestro D, Vigliotta G, Imperi F, Visca P, Alifano P, Parente D. 2007. Microbial community structure and dynamics of dark fire-cured tobacco fermentation. Appl Environ Microbiol 73:825–37. doi:10.1128/AEM.02378-06.

9. Huang J, Yang J, Duan Y, Gu W, Gong X, Zhe W, Su C, Zhang K-Q. 2010. Bacterial diversities on unaged and aging flue-cured tobacco leaves estimated by 16S rRNA sequence analysis. Appl Microbiol Biotechnol 88:553–62. doi:10.1007/s00253-010-2763-4.

10. Li J, Zhao Y, Qin Y, Shi H. 2020. Influence of microbiota and metabolites on the quality of tobacco during fermentation. BMC Microbiol 20. doi:10.1186/s12866-020-02035-8.

11. Zhou J, Yu L, Zhang J, Zhang X, Xue Y, Liu J, Xiao Z. 2020. Characterization of the core microbiome in tobacco leaves during aging. MicrobiologyOpen 9. doi:10.1002/mbo3.984.

12. Zhang Q, Geng Z, Li D, Ding Z. 2020. Characterization and discrimination of microbial community and co-occurrence patterns in fresh and strong flavor style flue-cured tobacco leaves. Microbiologyopen 9:e965. doi:10.1002/mbo3.965.

13. Tao J, Chen Q, Chen S, Lu P, Chen Y, Jin J, Li J, Xu Y, He W, Long T, Deng X, Yin H, Li Z, Fan J, Cao P. 2022. Metagenomic insight into the microbial degradation of organic compounds in fermented plant leaves. Environ Res 214:113902. doi:10.1016/j.envres.2022.113902.

14. Han F. 2010. Tobacco chemistry. China Agriculture Press.

15. Hur SJ, Lee SY, Kim YC, Choi I, Kim GB. 2014. Effect of fermentation on the antioxidant activity in plant-based foods. Food Chem 160:346–56. doi:10.1016/j.foodchem.2014.03.112.

16. Zhang C, Gao Z, Shi W, Li L, Tian R, Huang J, Lin R, Wang B, Zhou B. 2020. Material conversion, microbial community composition and metabolic functional succession during green soybean hull composting. Bioresour Technol 316:123823. doi:10.1016/j.biortech.2020.123823.

17. Frankenburg WG, Gottscho AM. 1955. The chemistry of tobacco fermentation. I. Conversion of the alkaloids. B. The formation of oxynicotine. J Am Chem Soc 77:5728–5730. doi:10.1021/ja01626a076.

18. Frankenburg WG, Gottscho AM, Mayaud EW, Tso T-C. 1952. The chemistry of tobacco fermentation. I. Conversion of the alkaloids. A. The formation of 3-pyridyl methyl ketone and of 2,3’-dipyridyl. J Am Chem Soc 74:4309–4314. doi:10.1021/ja01137a018.

19. Frankenburg WG, Gottscho AM, Vaitekunas AA, Zacharius RM. 1955. The chemistry of tobacco fermentation. I. Conversion of the alkaloids. C. The formation of 3-pyridyl propyl ketone, nicotinamide and N-methylnicotinamide. J Am Chem Soc 77:5730–5732. doi:10.1021/ja01626a077.

20. Frankenburg WG, Vaitekunas AA. 1957. The chemistry of tobacco fermentation. I. Conversion of the alkaloids. D. Identification of cotinine in fermented leaves. J Am Chem Soc 79:149–151. doi:10.1021/ja01558a039.

21. Shi X, Wang X, Lin K, Cui J, Li Z, Li L. 2013. Changes of aroma substances in cigar wrapper tobacco leaves during the stacking fermentation. Acta Agriculturae Boreali-Occidentalis Sinica 22:114–119. doi:10.16472/j.

22. Zheng T, Zhang Q, Li P, Wu X, Liu Y, Yang Z, Li D, Zhang J, Du G. 2022. Analysis of Microbial Community, Volatile Flavor Compounds, and Flavor of Cigar Tobacco Leaves From Different Regions. Front Microbiol 13:907270. doi:10.3389/fmicb.2022.907270.

23. Vigliotta G, Di Giacomo M, Carata E, Massardo DR, Tredici SM, Silvestro D, Paolino M, Pontieri P, Del Giudice L, Parente D, Alifano P. 2007. Nitrite metabolism in Debaryomyces hansenii TOB-Y7, a yeast strain involved in tobacco fermentation. Appl Microbiol Biotechnol 75:633–45. doi:10.1007/s00253-007-0867-2.

24. Dai J, Dong A, Xiong G, Liu Y, Hossain MS, Liu S, Gao N, Li S, Wang J, Qiu D. 2020. Production of highly active extracellular amylase and cellulase from bacillus subtilis ZIM3 and a recombinant strain with a potential application in tobacco fermentation. Frontiers in Microbiology 11:1539. doi:10.3389/fmicb.2020.01539.

25. Ge Z, Kaichao L, Yuhua X, Juan W, Ghugui L, Fang W, Haibo Z, Haobao L. 2018. Isolation and activity determination of surface bacteria in cigar wrapper leaves from four different countries. Chinese Tobacco Science 39:82–88. doi:10.13496/j.issn.1007-5119.2018.02.012.

26. Cecchini NM, Roychoudhry S, Speed DJ, Steffes K, Tambe A, Zodrow K, Konstantinoff K, Jung HW, Engle NL, Tschaplinski TJ, Greenberg JT. 2019. Underground Azelaic Acid-Conferred Resistance to Pseudomonas syringae in Arabidopsis. Mol Plant Microbe Interact 32:86–94. doi:10.1094/MPMI-07-18-0185-R.

27. Li Y, Lin q, Pang T, Shi J, Kong G, Mu x. 2015. Qualititative and quantitative analysis of volative flavor components in tobacco leaf by gas chromatography-mass spectrometry. Journal of Analytical Science 31:372–378. doi:10.13526/j.issn.1006-6144.2015.03.016.

28. Yu J, Gong C. 2009. Primary processing of tobacco raw materials. China Agriculture Press.

29. Zhu X, Gao Y, Chen Z, Su Q. 2009. Development of a chromatographic fingerprint of tobacco flavor by use of GC and GC-MS. Chromatographia 69:735–742. doi:10.1365/s10337-009-0968-4.

30. Haas BJ, Gevers D, Earl AM, Feldgarden M, Ward DV, Giannoukos G, Ciulla D, Tabbaa D, Highlander SK, Sodergren E. 2011. Chimeric 16S rRNA sequence formation and detection in Sanger and 454-pyrosequenced PCR amplicons. Genome Res 21:494–504. doi:10.1101/gr.112730.110.

31. Magoč T, Salzberg SL. 2011. FLASH: fast length adjustment of short reads to improve genome assemblies. Bioinformatics 27:2957–2963. doi:10.1093/bioinformatics/btr507.

32. Li M, Shao D, Zhou J, Gu J, Qin J, Chen W, Wei W. 2020. Signatures within esophageal microbiota with progression of esophageal squamous cell carcinoma. Chin J Cancer Res 32:755–767. doi:10.21147/j.issn.1000-9604.2020.06.09.

33. Aßhauer KP, Wemheuer B, Daniel R, Meinicke P. 2015. Tax4Fun: predicting functional profiles from metagenomic 16S rRNA data. Bioinformatics 31:2882–2884. doi:10.1093/bioinformatics/btv287.

34. Nguyen NH, Song Z, Bates ST, Branco S, Tedersoo L, Menke J, Schilling JS, Kennedy PG. 2016. FUNGuild: An open annotation tool for parsing fungal community datasets by ecological guild. Fungal Ecology 20:241–248. doi:10.1016/j.funeco.2015.06.006.

